# Gene Expression of Functionally-Related Genes Coevolves Across Fungal Species: Detecting Coevolution of Gene Expression Using Phylogenetic Comparative Methods

**DOI:** 10.1101/844472

**Authors:** Alexander L. Cope, Brian O’Meara, Michael A. Gilchrist

## Abstract

**Background:** Researchers often measure changes in gene expression across conditions to better understand the shared functional roles and regulatory mechanisms of different genes. Analogous to this is comparing gene expression across species, which can improve our understanding of the evolutionary processes shaping the evolution of both individual genes and functional pathways. One area of interest is determining genes showing signals of coevolution, which can also indicate potential functional similarity, analogous to co-expression analysis often performed across conditions for a single species. However, as with any trait, comparing gene expression across species can be confounded by the non-independence of species due to shared ancestry, making standard hypothesis testing inappropriate.

**Results:** We compared RNA-Seq data across 18 fungal species using a multivariate Brownian Motion phylogenetic comparative method (PCM), which allowed us to quantify coevolution between protein pairs while directly accounting for the shared ancestry of the species. Our work indicates proteins which physically-interact show stronger signals of coevolution than randomly-generated pairs. Interactions with stronger empirical and computational evidence also showing stronger signals of coevolution. We examined the effects of number of protein interactions and gene expression levels on coevolution, finding both factors are overall poor predictors of the strength of coevolution between a protein pair. Simulations further demonstrate the potential issues of analyzing gene expression coevolution without accounting for shared ancestry in a standard hypothesis testing framework. Furthermore, our simulations indicate the use of a randomly-generated null distribution as a means of determining statistical significance for detecting coevolving genes with phylogenetically-uncorrected correlations, as has previously been done, is less accurate than PCMs, although is a significant improvement over standard hypothesis testing. These methods are further improved by using a phylogenetically-corrected correlation metric.

**Conclusions:** Our work highlights potential benefits of using PCMs to detect gene expression coevolution from high-throughput omics scale data. This framework can be built upon to investigate other evolutionary hypotheses, such as changes in transcription regulatory mechanisms across species.

## Introduction

Analysis of high-throughput transcriptomics and proteomics data often focuses on how changes in environment (e.g. nutrient availability) result in changes in mRNA or protein abundances [1]. Through the concept of “guilt-by-association,” genes which show similar gene expression patterns across conditions are hypothesized to be functionally-related [2, 3, 4, 5]. For example, in *S. cerevisiae*, there is significant overlap between the proteins which physically interact and the proteins which are co-expressed [6]. Such observations have naturally led researchers to ask if functionally-related genes show coordinated changes in expression across conditions, do they also show coordinated changes, or coevolve, across species.

Previous work supports the hypothesis that gene expression of functionally-related genes shows stronger signals of coevolution than randomly-generated gene pairs in both unicellular yeasts and a diverse set of prokaryotes. [7, 8, 9]. Interestingly, the strength of this signal appeared to vary based on the functional groupings of the genes in question [7]. Fraser et. al. [8] proposed gene expression coevolution could be a useful method for predicting proteins which are functionally-related.

Most of the previous work examining coevolution of gene expression relied upon the Codon Adaptation Index (CAI) [10] as a proxy for gene expression. CAI and other codon-usage metrics often correlate well with gene expression in many species, but this is often not the case in species with a strong mutational bias or low effective population sizes, as is the case in many multicellular eukaryotes [11]. In fact, Lithwick and Margalit [9] were forced to eliminate organisms from their analysis which showed little adaptive codon usage. This makes detecting signals from empirical measures of gene expression, such as from RNA-Seq or mass spectrometry data, particularly useful for many species where codon usage metrics are a poor proxy for gene expression. Recent work by Martin and Fraser [12] demonstrated a method for examining coevolution of gene expression within sets of functionally-related genes using RNA-Seq data measured from the Marine Microbial Eukaryotic Transcriptome Project [13].

While it may seem appropriate to simply assess the correlation (e.g. Pearson or Spearman) between gene expression estimates across species, much like one might do in a co-expression analysis across conditions, an issue that arises is the non-independence of species due to shared ancestry [14]. This can result in biases in correlation coefficients and lead to an inflation of the degrees of freedom, making standard hypothesis testing inappropriate [14, 15]. Recent work concluded comparative analysis of gene expression data across species can be confounded by the phylogeny, leading potentially to incorrect inferences [16]. Previous work examining coevolution of gene expression did not directly account for the phylogeny when estimating correlation coefficients of gene expression across species, which is thought to reflect the strength of coevolution between gene pairs. With the exception of Clark et. al. [7], who applied a transformation to their correlation coefficients originally developed to eliminate phylogenetic signal from sequence coevolution data [17], much of the previous work used a randomly-generated null distribution created from genes not thought to coevolve as a means of determining a statistical significance cutoff. Although the use of a randomly-generated null is likely a better alternative than standard hypothesis testing, a direct assessment of these approaches’ abilities to adequately control for the phylogeny have not been determined, to the best of our knowledge.

An alternative solution is to directly account for the phylogeny when assessing coevolution between pairs of genes using phylogenetic comparative methods (PCMs). Previous efforts have developed PCMs for examining coevolution of functionally-related genes based on the presence/absence of genes across species. Barker and Pagel [18] developed what is essentially a phylogenetically-corrected version of phylogenetic profiling, which looks at the correlated presence/absence of genes across species. Looking across a set of fungal species and using protein-protein interaction data to determine functionally-related genes, they found incorporating the phylogeny reduced the false positive rate compared to a Fisher’s exact test. Of course, this method is not applicable if the genes are present in all species under consideration, making gene expression a valuable trait for investigating coevolution of functionally-related genes.

Many PCMs have been developed for studying the evolution of gene expression, although this work has not focused on detecting coevolution of gene expression. [19, 20, 21, 22, 23, 24, 25, 26, 27]. Much of this work relies on modeling gene expression evolution as an Ornstein-Uhlenbeck (OU) process [28, 29]. Modeling trait evolution as an OU process assumes the trait is evolving around an optimal value. A multivariate versions of the OU model exists [30], but the additional parameters used in the model often requires a greater amount of species-level data to make accurate parameter estimates. Here, we present an approach which models the coevolution of gene expression, as estimated via RNA-Seq, for pairs of proteins using the simpler multivariate Brownian Motion (BM) model [31, 32]. This approach allows us to estimate the degree of correlation between two traits over evolutionary time while accounting for the shared ancestry of the considered species.

We find physically-interacting proteins show, on average, stronger gene expression coevolution than randomly-generated pairs of proteins using the multivariate BM approach. We also find phylogenetically-uncorrected correlations tend to inflate estimates of gene expression coevolution. Unsurprisingly, simulations reveal standard hypothesis testing (i.e. *p* < 0.05) using phylogenetically-uncorrected correlations inflates the false discovery rate. We find determing statistical significance via a randomly-generated null distribution, as described in Fraser et. al. [8] is a significant improvement over standard hypothesis testing, but still performs worse than the PCM approach. The method recently described by Martin and Fraser [12] was able to obtain a low false discovery rate, but this came at the expense of statistical power to detect coevolving genes relative to the PCM, which had a comparable false discovery rate.

We expand upon previous work by looking for potential predictors reflecting the strength of coevolution between two pairs of proteins. As expected, we find protein pairs with stronger evidence of functional-relatedness tend show stronger coevolution at the gene expression level. We also find gene expression level and the number of protein interactions, which are considered good predictors of evolutionary rate of a gene [33], are poor predictors of the the strength of coevolution between protein pairs. Consistent with previous results, we also find coevolution of gene expression is an overall weak predictor of protein sequence coevolution.

## Materials and methods

### Protein Interaction Data

18 fungal species were chosen due to availability of RNA-Seq data and for comparability to previous studies examining the evolution of functionally-related proteins [8, 7, 18]. Consistent with [8] and [18], we use physically-interacting proteins as our test case for examining functionally-related proteins. The STRING database was used to identify empirically-determined protein-protein interactions in species for which data was available [34]. We assume these protein-protein interactions are conserved across all species under consideration. This dataset will be referred to as the “binding group” for the remainder of the paper. Randomly-generated protein pairs followed by removal of any pairs which were annotated in the STRING database for the species under consideration, even if the annotation did not specify a “binding” interaction. Any proteins with overlapping Gene Ontology terms were removed to control for potential false negatives. This dataset will be referred to as the “control group” for the remainder of the manuscript.

### Gene Expression Data

Gene expression levels were estimated from publicly available RNA-Seq datasets taken from SRA using the pseudo-alignment tool, Salmon [35]. Reads were mapped against protein-coding sequences obtained from NCBI Refseq, with the exception of *S. kudriavzevii*, *S. mikatae*, *S. paradoxus*, and *S. bayanus*, which were pulled from [36], and *L. kluyverii*, which was obtained from the Broad Institute (https://portals.broadinstitute.org/regev/orthogroups/) FASTQC was used to assess the quality of the RNA-Seq reads. If necessary, TrimGalore was used to remove adaptor sequences (https://www.bioinformatics.babraham.ac.uk/projects/trimgalore/). Gene expression counts were obtained using Salmon’s built-in ability to control for GC and position-specific biases, and these counts were converted to the transcripts per million (TPM) metric [37]. For single-end reads, mean and standard deviation for fragment lengths were specified to be 200 and 80, respectively, except for *S. mikatae*, *S. paradoxus*, *S. paradoxus*, for which mean fragment length was specified to be 250 [38].

Given the RNA-Seq experiments are often measured different conditions, we only selected samples from the control conditions, as these are more likely to reflect natural or standard conditions for a species. For datasets which were time course experiments, we randomly selected 3 time-points which were well-correlated in gene expression estimates (Pearson correlation *ρ* > 0.98). Each RNASeq sample/replicate for each species was transformed to a standard lognormal distribution (i.e. *ln*(*X*) *∼ N*(0, 1)), consistent with the transformation used by [19]. A mean and standard error of normalized TPM values were calculated for each gene across all samples/replicates used. Genes with missing data, which could be because no ortholog was identified between species or no gene expression estimate was obtained, were excluded from further analysis.

We note some of the RNA-Seq datasets did not indicate replicates, making it impossible to estimate a standard error measurement for the analysis. It is generally recommend measurement error be provided for the analysis of continuous traits during phylogenetic analysis. As a proxy for the species missing replicates, we used a closely-related species to provides estimates of the standard error. This included *S. paradoxus* (proxy: *S. cerevisiae*), *S. mikatae* (proxy: *S. bayanus*), and *N. tetrasperma* and *N. discreta* (proxy: *N. crasse*).

### Ortholog identification

Orthologs for fungal species were taken from FungiDB [39], previous publications [36, 40], or the Reciprocal Best Hits BLAST approach, which was only used for *N. castellii*. Proteins with an annotated paralog in the FungalDB or previous literature were excluded from the analysis, as introduction of a paralog could impact the gene expression of the original gene. This eliminated 4,307 possible genes.

### Phylogenetic tree construction

Codon alignments of 59 complete, randomly chosen nuclear ORF were performed using TranslatorX using the MAFFT option followed by GBlocks filtering to remove poorly aligned regions [41]. These alignments were concatenated, followed by phylogenetic tree estimation using RAxML with a partitioned GTR-Γ fit allowing rate parameters for the third codon position to vary from the first and second codon position. *C. neoformans* was designated as an outgroup. Under the Brownian Motion model, it is ideal to use an ultrametric tree unless extinct species are present [42]. To convert the RAxML tree to a ultrametric tree, treePL [43] was used to date the tree, taking the divergence time of *S. cerevisiae* and *C. neoformans* (723 MYA, from TimeTree [44]) as a calibration point. The final phylogenetic treee used for all analyses can be observed in Figure 1. A summary of the species used, the RNA-Seq data used, and the availability of protein-protein interaction data from STRING can be found in Additional File 1, Table S1.

**Figure 1:**
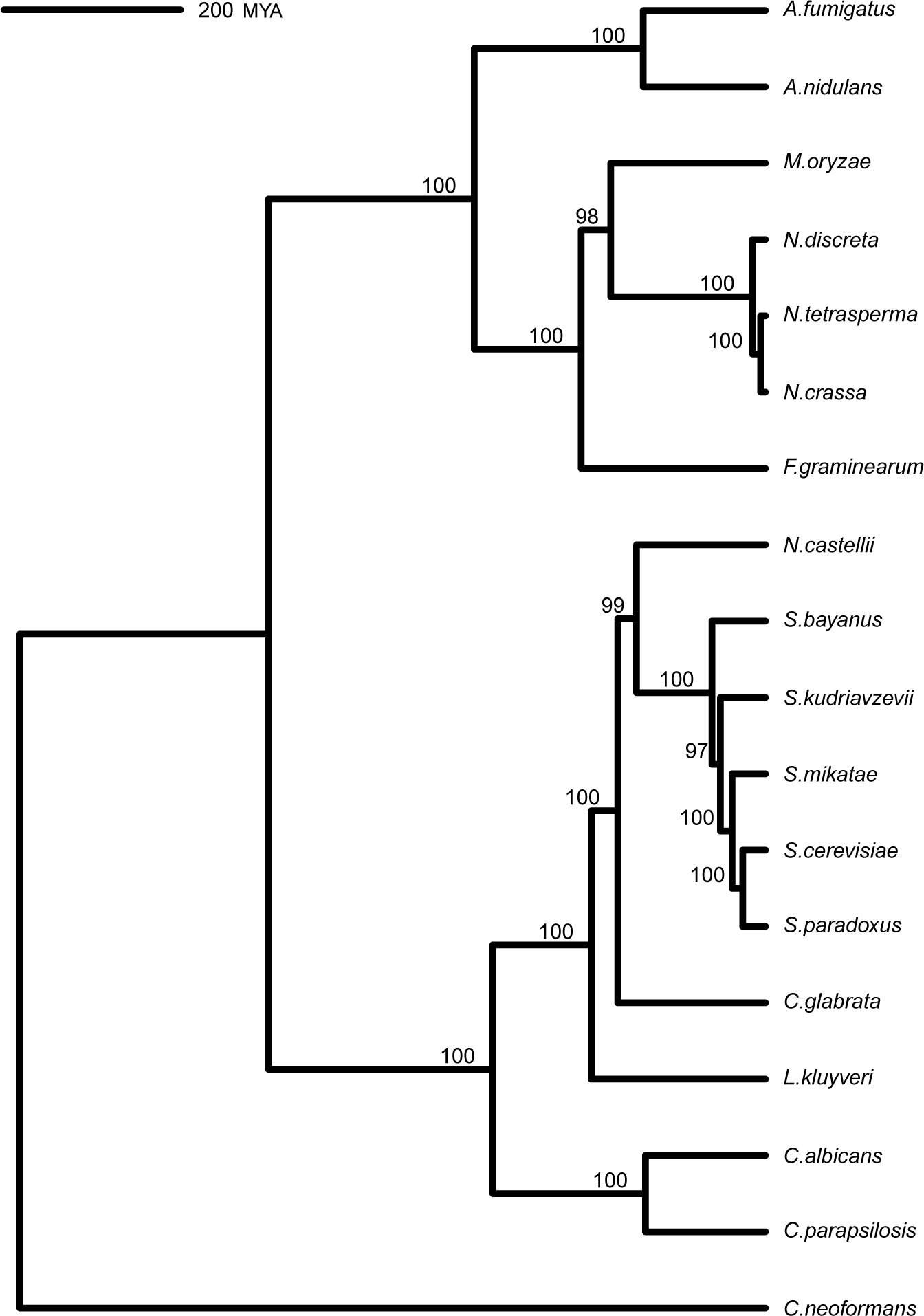
Dated phylogenetic tree with RAxML bootstrap support. Branch lengths are in millions of years.

### Analysis of Gene Expression Data

Analyses and visualizations were performed using the R programming language. Gene expression evolution was modeled as a multivariate Brownian Motion process using the R package **mvMORPH**. Briefly, the evolutionary rate matrix for multivariate Brownian Motion represents both the trait variances on the diagonal for the individual gene expression values, as well as the trait covariance between the gene expression estimates on the off-diagonal. The evolutionary correlation coefficient *ρ*_*C*_ reflects the degree to which gene expression estimates are correlated over evolutionary time and can be calculated from the evolutionary rate matrix [31, 32, 45]. The evolutionary correlation coefficient *ρ*_*C*_ will from here on out be referred to as the “phylogenetically-corrected correlation” to emphasize this statistic accounts for the shared ancestry of the species. Likewise, we will refer to the Pearson correlation coefficient *ρ*_*U*_ (estimated via the R built-in function cor.test()) as the “phylogenetically-uncorrected correlation”, as this statistic ignores shared ancestry and uses variances and covariances estimated from the data at the tips of the tree.

Appropriateness of the Brownian Motion for modeling trait evolution was assessed as described in [46]. Briefly, phylogenetic independent contrasts (PICs) and standardized variances [14] were calculated from gene expression data for each ortholog set using the pic() function from the **ape** R package [47]. Pairs of genes containing a significant correlation (i.e. *p* < 0.05) between PICs and standardized variances, which indicates violation of Brownian Motion assumptions [42, 46], were excluded from further analyses.

The phylogenetically-corrected correlation *ρ*_*C*_, which reflects the strength of gene expression coevolution between two genes, was compared to metrics associated with functional-relatedness of two genes. We expect stronger coevolution of gene expression between proteins which are more functionally-related. As a metric of functional-relatedness for each interaction, we used the STRING confidence score, which factors in both empirical/computational evidence supporting an interaction, as well as evidence from closely-related species. Similarly, one might expect proteins sharing a greater number of overlapping Gene Ontology (GO) terms to be more functionally-related.

It is well-established both gene expression and number of interactions in a protein-protein interaction network impact the evolutionary behavior of a protein [48, 49]; thus, we also tested if such protein-level properties also impact the strength of coevolution between two proteins. We hypothesized proteins pairs which are, on average, more highly expressed and involved in more interactions would show stronger coevolution of gene expression. For each protein pair in the binding group, the mean degree (i.e. the average number of interactions for each protein) and the mean phylogenetically-corrected average gene expression value were calculated. The phylogenetically-corrected average gene expression value for a protein is taken as the ancestral state value estimated at the root of the tree by **mvMORPH**.

To determine if functional-relatedness, gene expression, and number of protein interactions have an impact on the strength of coevolution, a weighted rank-based (i.e. robust to non-normality in data) Spearman correlation *ρ*_*S*_ was used to reduce the impact of proteins found in multiple pairs. Weights for the weighted Spearman correlation *ρ*_*S*_ for each protein pair were calculated as

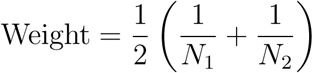

where *N*_*i*_ is the the number of times protein *i* appears in the binding group. Confidence intervals and p-values for the weighted Spearman correlations were calculated using the R package **boot** [50, 51].

To assess the impact of proteins found in multiple pairs on differences observed between the binding and control groups, we generated 200 subsets of the binding and control datasets in which a protein was only allowed to appear, at maximum, in one protein pair per dataset. Each subset was restricted to a maximum size of 200 protein pairs. For each subset, the mean was calculated for *ρ*_*C*_ and *ρ*_*U*_, creating a distribution of means. Scripts and for performing phylogenetic analysis and post-analysis of the results can be found at https://github.com/acope3/GeneExpression_coevolution.

### Assessing accuracy of methods for detecting coevolution of gene expression

Data simulated under Brownian Motion were used to assess the ability to detect coevolution of gene expression (see Additional File 1 for details). For the PCM approach, protein pairs were considered coevolving if a Likelihood Ratio test (as implemented in **mvMORPH**) comparing the model allowing coevolution of gene expression to a null model forcing independent evolution of gene expression had a Benjamini-Hochberg corrected p-value *<* 0.05. Similarly, for the non-PCM approach (cor.test() function in R), protein pairs were considered significantly coevolving if the phylogenetically-uncorrected correlation *ρ*_*U*_ had a Benjamini-Hochberg corrected p-value *<* 0.05

Previous work proposed using randomly-generated null distributions (i.e. the control group) as a means of determining statistically significant gene expression coevolution using phylogenetically-uncorrected correlations. This approach is thought to be an adequate approach to control for the phylogeny when the phylogeny is unknown [12]. We implement approaches similar to those described in Fraser et. al. [8] and Martin and Fraser [12] using both the phylogenetically-uncorrected and phylogenetically-corrected correlations.

Fraser et. al. compared the relative histograms of correlations from a binding and a control group to determine the bin at which the relative frequencies of the binding group were greater than the control group for all subsequent bins. Pairs of proteins were considered significantly coevolving if they had a correlation greater than this point. To assess the accuracy of this method, we split both the binding and control groups into training and test sets (80% and 20% of the data, respectively). The binding and control training sets were used to determine the significance cutoff, while the test sets were then used to assess the accuracy of this approach.

Martin and Fraser presented an approach to determine if gene sets (i.e. more than 2 genes) showed significant coevolution of gene expression by comparing the median phylogenetically-uncorrected correlation to the median correlations from 10,000 randomly-generated gene sets. As we only deal with protein pairs, we compared the number of times (out of 1000) a randomly-generated protein pair had a correlation greater than the correlation of the target protein pair. This procedure was repeated for each protein pair in the binding and control groups. A p-value for each pair was calculated as described in Martin and Fraser [12], and a p-value cutoff was empirically-determined such that the false discovery rate was approximately 5%.

We note accuracy scores can be skewed by large differences in the size of the binding and control groups. For example, if a method is underpowered and the size of the control group is much larger than the binding group, then failure to detect significant differences in the binding group is heavily outweighed by successfully not detecting significant differences in the control group. This results in a higher, and potentially misleading, accuracy score for the method. To account for this, each method was assessed using a subsample of the control group which is the same size as the binding group. Model assessments were made 100 times to obtain mean true positive rates, false positive rates, false discovery rates, and overall accuracy scores.

## Results

Overall, the normalized gene expression data are moderately to strongly correlated between all species (Additional File 1, Figure S1). Clearly, species which are more closely-related tend to show stronger correlations between normalized gene expression values, consistent with expectations. The *Candida* species appear to be exceptions, but these yeast demonstrate pathogenic traits, which could partially explain some of these differences, as well as why two of these species (*C. glabrata* and *C. parapsilosis*) appear to be better correlated with the pathogenic *Aspergillus* species.

After filtering proteins based on missing data or violation of the Brownian Motion assumption, our binding and control datasets contained 3,091 and 13,936 protein pairs respectively, consisting of 648 unique proteins. We note similar patterns are observed if not excluding genes which violate the BM assumption, although the signal appears weaker (see Additional File 1, Figure S6 – S9).

### Interacting proteins demonstrate clear coevolution of gene expression

Examination of the phylogenetically-corrected correlation *ρ_C_* distribution for the binding and control group shows striking differences (Figure 2) Binding proteins have a mean phylogenetically-corrected correlation of 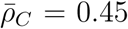 (One-sample t-test, 95% CI: 0.436 – 0.464, *p* < 10^*−*200^). In contrast, the randomly-generated control group had a much lower mean phylogenetically-corrected correlation of 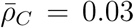 (One-sample t-test, 95% CI: 0.025 – 0.037, *p <* 10^*−*23^). As is clear from the 95% confidence intervals, the difference between the mean phylogenetically-corrected correlations for the binding and control distributions is statistically significant (Welch’s t-test, *p <* 10^*−*200^). Although the mean phylogenetically-corrected correlation for the control group is significantly different from 0, it is important to note two things: (1) even though we did our best to eliminate possible false negatives in the control group, it is unlikely all false negatives were eliminated and (2) this is consistent with previous work by [8], who also had random control groups which were not centered around 0. Despite the small, but statistically significant, deviation from 0 of the control group, the binding group shows a clear skew towards stronger coevolution between protein pairs than is observed in the control group, as expected.

**Figure 2:**
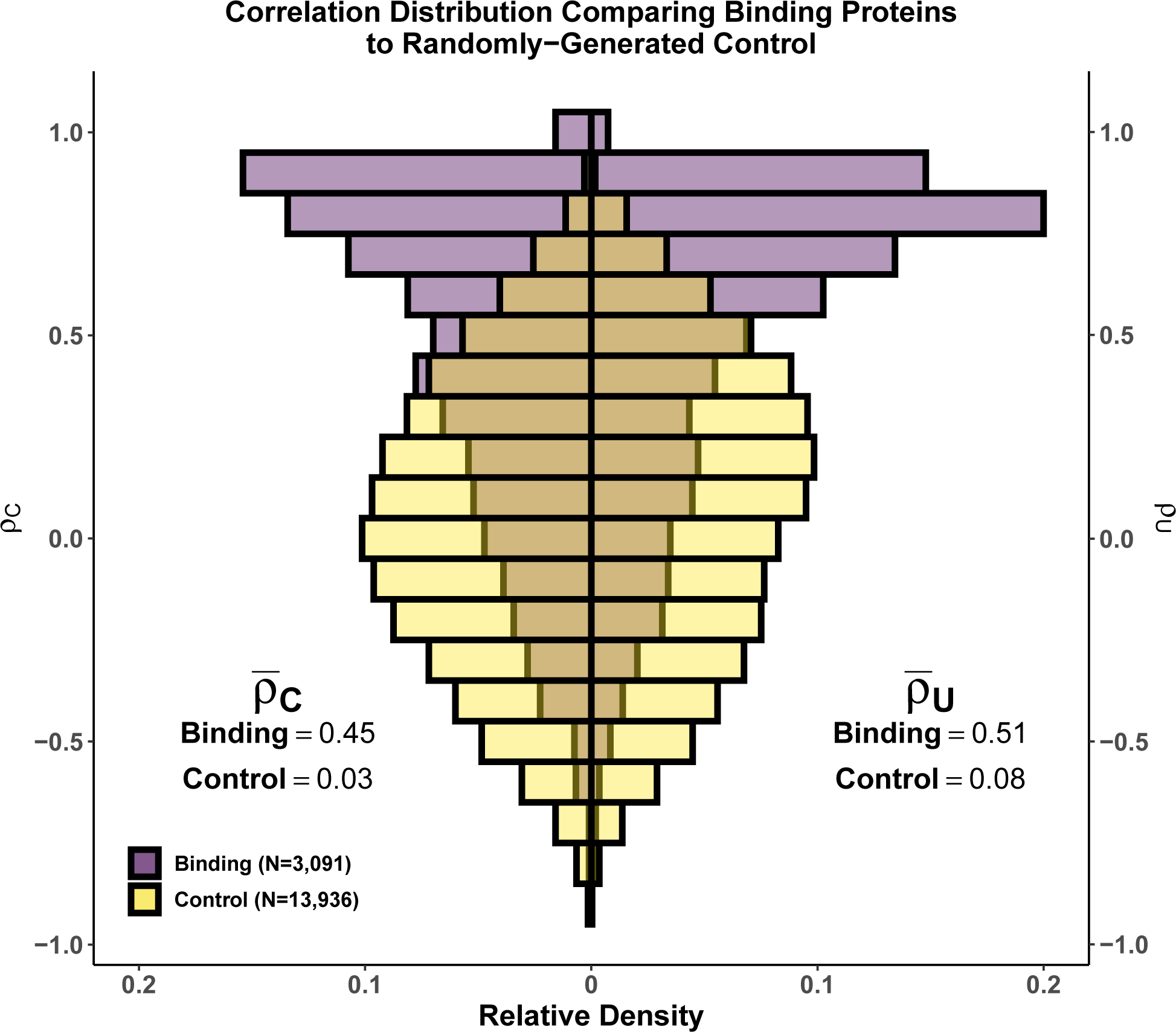
Comparing phylogenetically-corrected and uncorrected correlations. Comparing the distributions of the (Left) phylogenetically-corrected correlation *ρ*_*C*_ and the (Right) phylogenetically-uncorrected correlation *ρ*_*U*_ for the binding (purple) and control (yellow) groups. (Left) Mean values for the binding and control group phylogenetically-corrected correlation *ρ*_*C*_ distributions are 0.45 (95% CI: 0.436 – 0.464) and 0.03 (95% CI: 0.025 – 0.037), respectively. (Right) Mean values for the binding and control group phylogenetically-uncorrected correlation *ρ*_*U*_ distributions are 0.51 (95% CI: 0.497 – 0.523) and 0.08 (95% CI: 0.074 – 0.086), respectively

We find a weak, but significant, positive correlation between the STRING confidence scores and phylogenetically-corrected correlations *ρ*_*C*_ (Figure 3, Weighted Spearman Rank Correlation *ρ*_*S*_ = 0.32, 95% CI: 0.274 – 0.371, *p <* 10^*−*37^), indicating interactions which are more likely to be true and conserved show stronger coevolution of gene expression (Figure 3). A similar result is obtained when using a metric of functional similarity between proteins based on overlapping Gene Ontology terms (Additional File 1, Figure S2).

**Figure 3:**
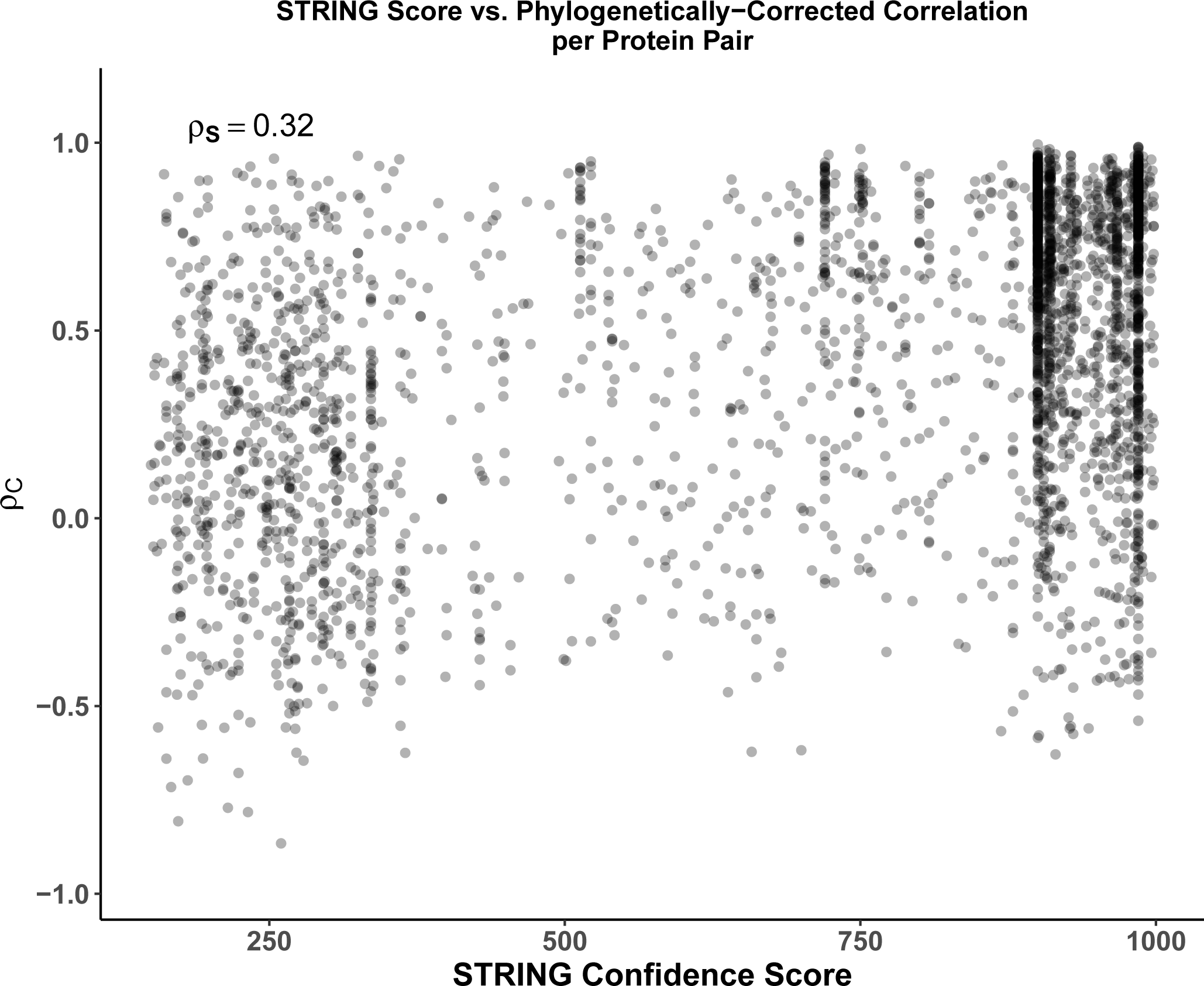
Effects of functional-relatedness on phylogenetically-corrected correlation *ρ*_*C*_. Positive weighted Spearman rank correlation (*ρ*_*S*_ = 0.32, *p <* 10^*−*37^) between the STRING score and phylogenetically-corrected correlation *ρ*_*C*_ indicates more confident and/or conserved interactions tend to have higher *ρ*_*C*_, indicating stronger coevolution at the gene expression level.

We also compared how our phylogentically-corrected approach worked compared to a phylogenetically-uncorrected approach (Figure 2). Qualitatively, a similar pattern to the phylogenetically-corrected correlations *ρ*_*U*_ is observed: binding proteins show correlations positively skewed away from 0, consistent with stronger coevolution of gene expression between the interacting pairs. Interacting proteins had a mean phylogenetically-uncorrected correlation of 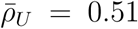 (One-sample t-test,95% CI: 0.497 – 0.523, *p <* 10^*−*200^). In contrast, randomly-generated protein pairs had a mean phylogenetically-uncorrected correlation 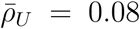 (One-sample t-test, 95% CI: 0.074 – 0.086, *p <* 10^*−*141^). As with the phylogenetically-corrected correlations, the control group deviates significantly from the null expectation of 0.0; however, the phylogenetically-uncorrected correlation deviates further from the expectation than the phylogenetically-corrected correlations. This is consistent with potential biasing of correlation estimates due to treatment of non-independent species data as independent [14, 15].

Simulations were performed to confirm potential problems with the use of non-phylogenetic methods for comparing gene expression across species (see Additional File 1). Results show failure to account for the phylogeny on data simulated under the null hypothesis of no coevolution between gene expression results in an increase in the false discovery rate (FDR, Table 1), consistent with expectations. However, the distribution of *ρ*_*U*_ simulated under no coevolution differs from the distribution of *ρ*_*U*_ from the real data (Additional File 1, Figure S5). In the case of simulated data in which no coevolution was allowed, the distribution of phylogenetically-uncorrected correlations *ρ*_*U*_ is centered around 0.0, unlike in the real data, but shows a broadening of the distribution compared to the phylogenetically-corrected correlations *ρ*_*C*_.

**Table 1:**
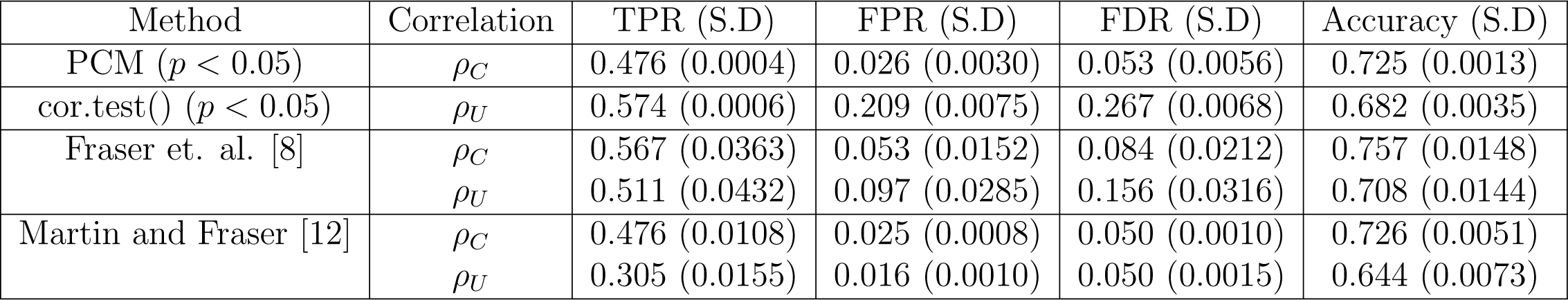
Mean and standard deviations of performance metrics for detecting coevolution of gene expression using simulated data. While standard hypothesis testing based on the phylogenetically-uncorrected correlation *ρ*_*U*_ (i.e. cor.test(), Benjamini-Hochberg corrected *p <* 0.05) has a greater true positive rate (TPR) compared to the PCM, it also has a much higher false discovery rate (FDR) when the true model is Brownian Motion. This is somewhat offset by using the method described in Fraser et. al. [8], but this does not completely control for the phylogeny as evidenced by a still high FDR (0.156). The method described by Martin and Fraser [12] can obtain an FDR of 0.05, but this drastically lowers its true positive rate if gene expression is evolving according to Brownian Motion. Both methods are improved by using the phylogenetically-corrected correlation *ρ*_*C*_.

Instead of determining statistical significance for the phylogenetically-uncorrected correlations *ρ*_*U*_ using *p <* 0.05, we used approaches similar to those described by Fraser et. al [8] and Martin and Fraser [12]. We found the method described in Fraser et. al. to have a greater true positive rate (TPR) compared to the PCM (0.511 compared to 0.476), but still had an inflated false discovery rate (FDR) of 0.156, although this was a significant improvement over standard hypothesis testing (Table 1). An approach similar to Martin and Fraser [12] was actually underpowered compared to the PCM, with a true positive rate (TPR) of 0.305, when controlling the FDR to be 0.05. This method had the overall worst accuracy of 0.644. Unsurprisingly, both methods described by Fraser et. al. and Martin and Fraser are improved when using the phylogenetically-corrected correlation *ρ*_*C*_. When the data contains phylogenetic signal, methods based on *ρ*_*C*_ are superior to the methods based on *ρ*_*C*_.

We note these methods all have fairly low true positive rates (TPR). We hypothesized part of this could be due to the presence of false positives in the binding group, which are unlikely to show much coevolution of gene expression, resulting in protein pairs in the simulated data with potentially small effects unlikely to be detected with only 18 species. After excluding potential false positives in the binding group (i.e. protein pairs with a STRING Score *<* 400), the TPR and overall accuracy of all methods increased (Additional File 1, S2). However, the general pattern remained the same: if the data contains phylogenetic signal, then methods based on correcting for the phylogeny are superior.

### Gene expression and number of interactions are poor predictors of coevolution of gene expression

It is well-established both gene expression and location in a protein-protein interaction network significantly impact the evolutionary behavior of a protein [49, 52, 53, 54, 55]. One might expect an imbalance in the number of proteins involved in a greater number of interactions or more highly expressed interactions to have a more negative impact on fitness, leading to greater constraints on the evolution of gene expression. However, we find both the number of interactions and the gene expression to be weak predictors of the strength of coevolution of gene expression. Based on the number of interactions for each protein in our binding dataset, the weighted Spearman rank correlation between the number of interactions and the phylogenetically-corrected correlations *ρ*_*C*_ is *ρ*_*S*_ = 0.26 (Figure 4a, 95% CI: 0.196 – 0.315, *p <* 10^*−*16^), indicating protein pairs involved in more interactions tend to show stronger constraint on the evolution of gene expression. Surprisingly, the mean ancestral gene expression estimates are negatively correlated with the phylogenetically-corrected correlations *ρ*_*C*_, with *ρ*_*S*_ = *−*0.09 (Figure **??**, 95% CI: - 0.143 – −0.035, *p* = 0.00131).

**Figure 4:**
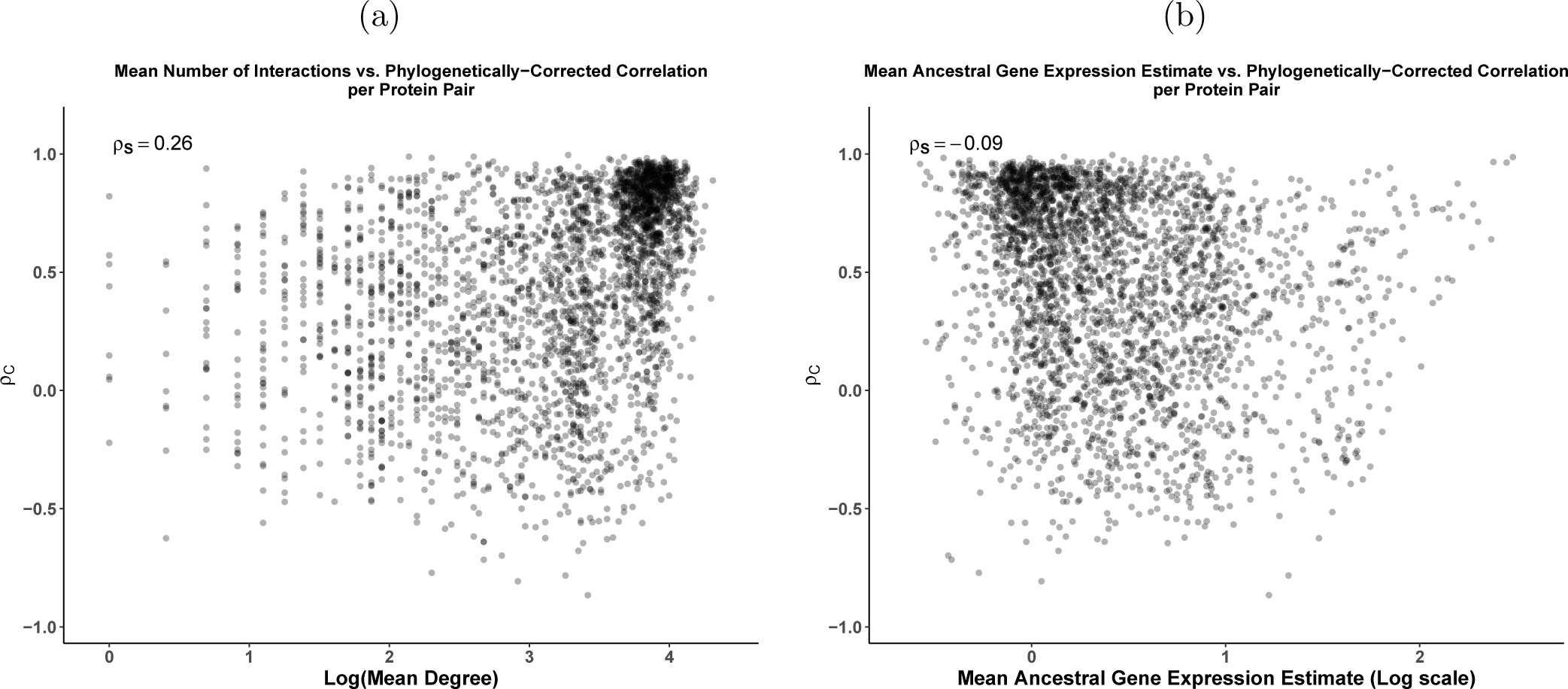
The relationship of (a) the mean degree (average number of interactions between a protein pair) and (b) mean ancestral gene expression estimate with the phylogenetically-corrected correlation *ρ*_*C*_ for the binding group. Both protein pair metrics are weakly, but significantly correlated with the phylogenetically-corrected correlation *ρ*_*C*_: weighted Spearman rank correlation *ρ*_*S*_ = 0.26 (*p <* 10^*−*16^) for mean degree and *ρ*_*S*_ = −0.09 (*p* = 0.00131) for mean ancestral gene expression. This suggests both metrics are poor predictors of the strength of coevolution of gene expression between protein pairs.

Given phylogenetically-corrected correlations *ρ*_*C*_ correlate with the number of interactions and mean ancestral gene expression, differences between the binding and control groups in terms of number of interactions and gene expression could introduce small biases when comparing the *ρ*_*C*_ distributions. The average mean ancestral gene expression estimate distributions for the binding and control group are extremely similar (0.414 vs. 0.416, respectively, Welch’s t-test, *p* = 0.8316). This makes differences in the gene expression distributions an unlikely source of bias when comparing the binding and control groups. To determine if protein membership biases causes biases in the results, 200 subsets of the binding and control groups were sampled, restricting a protein appearing in each group a maximum of 1 time. The 200 subsets resulted in distributions of the mean phylogenetically-corrected correlations 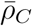, which were qualitatively consistent with the full datasets. We do note there appears to be less of a difference between the binding and control group 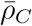 distributions compared to 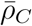 estimated from the full dataset (Additional File 1, Figure S3). This could be due to the representation of certain proteins in the binding group inflating the correlation, or could be due to decreased power to detect differences due to the significantly reduced dataset. Despite this, the overall interpretation is the same: interacting proteins show greater coevolution at the gene expression level than randomly generated pairs of proteins.

### Coevolution of gene expression weakly reflects coevolution of protein sequences

Previous work found an overall weak correlation between coevolution at the protein sequence level and coevolution at the gene expression level based on CAI [7, 8]. Using estimates of protein sequence coevolution across a yeast phylogeny taken from Clark et. al. [7], we found protein sequence coevolution and the phylogenetically-corrected correlations *ρ*_*C*_ were weakly, but significantly correlated (Weighted Spearman Rank correlation *ρ*_*S*_ = 0.10, 95% CI: 0.037 – 0.155, *p* = 0.0015, 5a). We also found a significant correlation between our phylogenetically-corrected correlation *ρ*_*C*_ and the measure of gene expression coevolution from Clark et. al. [7] (Weighted Spearman Rank correlation *ρ*_*S*_ = 0.22, 95% CI: 0.171 – 0.275, *p <* 10^*−*16^, Figure 5b). We find overall better agreement between CAI and empirical-based measures of coevolution for protein pairs which are, on average, more highly expressed (Weighted Spearman Rank correlation *ρ*_*S*_ = *−*0.12, 95% CI: −0.176 – −0.065, *p <* 10^*−*4^, Additional File 1, Figure S4). This is unsurprising, given that many highly expressed genes are likely to be housekeeping genes, such as ribosomal proteins, and thus highly expressed across most conditions and evolutionary time, making CAI a reliable proxy for gene expression in these cases.

**Figure 5:**
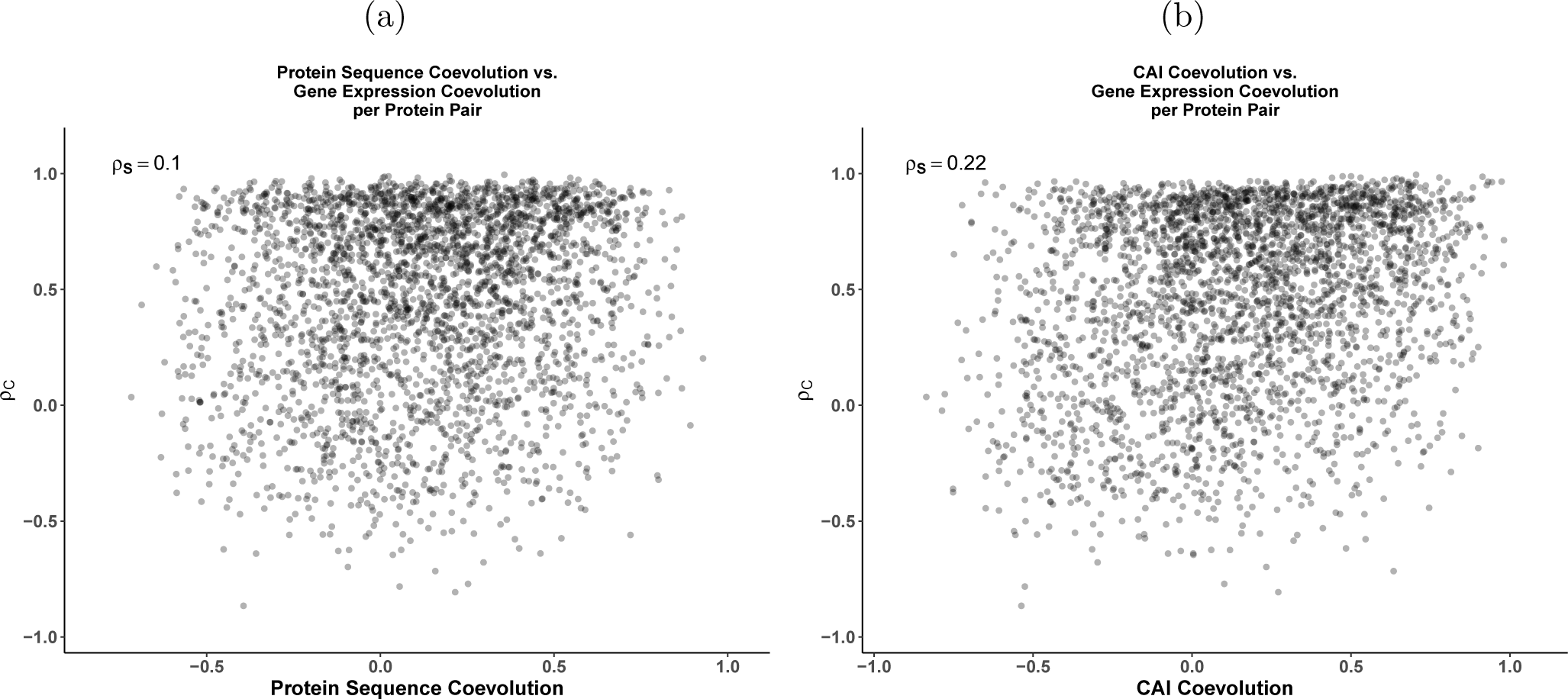
(a) Comparing coevolution of gene expression, represented by the phylogenetically-corrected correlation *ρ*_*C*_, and protein sequences, taken from Clark et al [7]. There is a weak but significant correlation (Weighted Spearman Rank Correlation *ρ*_*S*_ = 0.10, *p* = 0.0015) between the measures of gene expressions and protein sequence coevolution. (b) A similar comparison using the measures of CAI coevolution from [7]. Again, there is a weak, but significant correlation (Weighted Spearman Rank correlation *ρ*_*S*_ = 0.22, *p <* 10^*−*16^).

## Discussion

Consistent with previous results, we find physically-interacting proteins show a clear signal of gene expression coevolution compared to randomly-generated pairs of proteins, with mean phylogenetically-corrected correlations 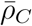 of 0.45 vs. 0.03, respectively. We find interacting proteins are correlated with the STRING confidence score (weighted Spearman Rank correlation *ρ*_*S*_ = 0.32), indicating protein-protein interactions with stronger evidence of being true and conserved have stronger co-evolution of gene expression, on average. We also find the number of protein-protein interactions a protein is involved in and its gene expression level – two common metrics known to affect the evolution of protein sequence – are overall weak predictors of gene expression coevolution. Protein pairs involved in more interactions do tend to show stronger gene expression coevolution (weighted Spearman rank correlations *ρ*_*S*_ = 0.26), consistent with the idea that proteins involved in more interactions in a protein-protein interaction network have more constraints on the evolution of their gene expression. Surprisingly, highly expressed protein pairs actually tended to show weaker coevolution of gene expression (weighted Spearman rank correlation *ρ*_*S*_ = *−*0.09). We also find an overall weak correlation between gene expression coevolution and protein sequence coevolution (weighted Spearman rank correlation *ρ*_*S*_ = 0.10), consistent with previous work [7, 8]. This is likely because relatively small regions of two protein sequences may be important for the proteins to be able to bind, forcing strong sequence coevolution at the binding sites, but weaker coevolution for the remainder of the protein sequences.

Surprisingly, there was overall poor agreement between CAI coevolution from Clark et. al. [7] and our measure of of gene expression coevolution based on empirical RNA-Seq data (weighted Spearman Rank correlation *ρ*_*S*_ = 0.22). The stronger correlation between *ρ*_*C*_ and CAI coevolution compared to protein sequence coevolution is unsurprising. CAI and similar codon usage metrics often show moderate to strong correlations with empirical gene expression estimates. However, the correlation between *ρ*_*C*_ and CAI coevolution is still very weak, indicating these measures of gene expression coevolution can give radically different interpretations about the degree of gene expression coevolution at the individual protein-pair level. It is worth noting that our estimates of gene expression coevolution and the estimates from [7] do not come from the same 18 species. Clark et. al. [7] also used 18 fungal species, 11 of which are from the *Saccharmoyces* or *Candida* genera, of which 7 overlap with the species used in this study. This undoubtedly introduced noise into these comparisons. Additionally, empirical gene expression in inherently noisy and can vary between conditions, which also will also increase the discrepancy between CAI and gene expression, particularly for low to moderate expression genes. Fortunately, many PCMs allow for the incorporation of measurement error, and through the use of multivariate methods, gene expression estimates across conditions can also be compared by treating each condition as an additional trait.

Unlike previous approaches, our results are based on both a multivariate PCM and empirical gene expression data. This offers two clear advantages. One advantage is our approach directly accounts for the phylogeny, recognizing the non-independence of species, allowing for standard hypothesis testing. Although previous efforts attempted to control for the phylogeny by using randomly-generated null distributions to determine statistical significance for phylogenetically-uncorrected correlations, our simulations indicate these approaches are generally worse than phylogenetic-based approaches <u>if</u> gene expression data contains phylogenetic signal (Table 1). The second advantage is while CAI often correlates well with gene expression in organisms with a high effective population size [11], low effective population size species often show little adaptive codon usage bias, making CAI a poor proxy for gene expression. As a result, the use of empirical gene expression measurements are highly valuable for studying the evolution of gene expression, as others have noted [1].

Our results indicate this multivariate PCM could be used to identify functionally-related proteins. However, simulations indicate more species might be needed to have sufficient statistical power (see Table 1), although this could vary depending on the tree and data in question. In theory, it is possible to expand this approach to test for gene expression coevolution in larger gene sets or correlate changes in gene expression with changes in other phenotypes, such as body size (see [45] for more details on using **mvMORPH**). With that in mind, recent work finds multivariate PCMs are in need of improvement, as parameter estimation accuracy decreases quickly as the number of traits (i.e. parameters) increases [56]. For now, it appears best to restrict the analysis to as few traits as possible. This is a case where the method developed by Martin and Fraser [12], which is designed to detect coevolution of gene expression in gene sets containing more than two genes, could be useful.

We note very few traits in biology likely evolve in a true Brownian Motion manner [14]. Consistent with this, most of the genes in our dataset violated the BM assumption based on the test proposed by [42]. Although the Ornstein-Uhlenbeck (OU) model may be a more appropriate model, and is used in many other PCMs for examining gene expression evolution, it often requires more species to make accurate parameter estimates. As we only used 18 fungal species, we opted to use the simpler Brownian Motion model combined with filtering of genes which significantly deviated from assumptions of the Brownian Motion model [42]. Future work should focus on the examination of coevolution of gene expression using the OU model. Based on our results, inclusion of genes which violate the BM assumption does not change overall conclusions of this work, but it does appear to weaken some of the observed signals (Additional File 1 Figure S6 – S9). Other phylogenetic models, such as the OU model, may be more appropriate for the analysis of gene expression for other gene pairs. A major advantage of PCMs is other models can easily be incorporated into the analysis of the trait, with the best model being determined via a hypothesis testing (e.g. Likelihood ratio test) or model comparison (e.g. AIC) framework.

We also note comparison of RNA-Seq data across species presents its own challenges [1, 37, 57]. Other methods for normalizing RNA-Seq data in order to make them comparable across species have been proposed, but to the best of our knowledge, there is no current consensus on the best approach. Brawand et. al. [20] developed a method for normalizing gene expression by identifying the genes with the most conserved ranks across samples, calculating species-specific scaling factors to make the median expression of these conserved rank genes equal across all species, and using those scaling factors to re-scale all gene expression estimates. Dunn et. al. [1] proposed a method based on comparing fold-changes (differential expression) across species-specific samples, which assumes a clear control and experimental condition and these measurements exists for all species under consideration. Muesser and Wagner [57] proposed a method for re-scaling the TPM metric based on the largest genome in the dataset, but this assumes the genes represented in the smaller genomes are subsets of the genes in the larger genome, which was not the case for our data based on the orthologs we identified. We transformed species-level data to a standard lognormal distribution, consistent with previous work using microarray data [19]. We make no claims that this is the best normalization approach for across species comparisons of RNA-Seq data, but state it was suitable for our overall purpose of determining if functionally-related genes show stronger coevolution of gene expression than randomly-generated pairs. Additionally, the log-transformed data likely better reflects the assumptions of Brownian Motion, as this eliminates the lower bound of 0.0 [42].

The RNA-Seq data used in this study were pulled from various non-related experiments which differed in terms of protocols, sequencers, sequencing depth, read type (single vs. paired), experimental conditions, and other factors which could impact the quantifications. It cannot be understated that this also introduces large amounts of variability to the quantified RNA-Seq data, making comparisons across species even more difficult. We attempted to control for this by using Salmon’s abilities to automatically adjust quantifications based on biases its detects within the RNA-Seq reads, as well as using the control conditions for each species for our analysis. Undoubtedly, this did not control for all of the variability introduced by pulling data from different experiments. Despite this, the normalized gene expression data were moderately to strongly correlated across species (Additional File 1, Figure S1) and species which were more closely related tended to show higher correlations, consistent with expectations. However, analyses attempting to make more precise conclusions about the evolution or coevolution of gene expression should ideally use measurements produced under better controlled conditions. Additionally, future efforts in this area may consider using proteomics data as opposed to transcriptomics. Previous work finds protein abundances appear to be more conserved between species compared to mRNA abundances, which could indicate stronger selection on maintaining the former [58].

Given our results and the ease of use of many tools implementing PCMs, we strongly recommend the use of PCM approaches when performing interspecies analysis. The phylogenetic research community has databases where phylogenetic trees can be easily accessed, such as TreeBase [59]. If a phylogenetic tree is not available for the species of interest, multiple sequence alignment tools and phylogenetic tree estimation tools have made building a reasonable phylogenetic tree efficient and easy, even for non-computational researchers. The phylogenetics community has made access to complex phylogenetic parameter estimation accessible via open-source, easy-to-use R packages, such as **mvMORPH** [45]. Although we strongly recommend the use of PCMs for interspecies data analysis, we emphasize that such approaches come with their own challenges and, in some cases, the PCM may not perform better than standard statistical approaches (see [46] for more details). Even so, approaches for assessing the impact of shared ancestry on the data still requires the generation of a phylogenetic tree and analysis of the trait in a phylogenetic context. Rohlfs et. al. also suggested PCMs likely will not provide different results from non-PCMs if analyzing gene expression for a small number of species, with a larger number of species resulting in more complex phylogenetic patterns and complicating the downstream data analyses [26]. Researchers should assess the impact of phylogeny of their data and make the appropriate decisions on what tools best answer the questions at hand.

## Declarations

### Ethics approval and consent to participate

Not applicable.

### Consent for publication

Not applicable.

### Availability of data and materials

References for all publicly-available sequencing data used for estimation of gene expression in this paper can be found in Additional File 1, Table S1. This includes references to the published manuscripts. Scripts for running the analysis described in this paper can be found on GitHub, as described in the Materials and Methods section.

### Authors’ contributions

ALC performed all analyses described in this work. BCO and MAG assisted in interpretation of data and writing of the manuscript. All the authors read and approved the final manuscript.

## Acknowledgements

Project was originally developed by A.L Cope and B.C. O’Meara as part of the NSF-funded Phylogenetic Methods course (NSF BIO 1453424) taught at the University of Tennessee, Knoxville. Financial support provided to A.L. Cope by NSF grant MCB-1546402 (A. VonArnim and M.A. Gilchrist), NSF grant MCB-1120370 (M.A. Gilchrist), DEB-1355033 (B.C. O’Meara, M.A. Gilchrist, and R. Zaretzki), the Graduate School of Genome Science and Technology (University of Tennessee), and the U.S. Department of Energy, Biological and Environmental program through funding of the Center for Bioenergy Innovation at the Oak Ridge National Laboratory. ORNL is managed by the UT – Battelle, LLC for the U.S. Department of Energy (DOE). Additional support was provided by the National Institute for Mathematical and Biological Synthesis (NSF:DBI-1300426) and the Dept. of Eco/Evol. Biology (University of Tennessee).

## Additional Files

**Additional File 1. Supplemental Material and Methods. Files contains supplemental materials and methods, including supplementary tables and figures referenced in this manuscript**.

