## Supplemental Information for "Gene Expression of Functionally-Related Genes Coevolves Across Fungal Species: Detecting Coevolution of Gene Expression Using Phylogenetic Comparative Methods"

### Supplemental Information: Gene Expression of Functionally-Related Genes Co-evolves Across Fungal Species

### Supplemental Methods

| Species | Phylum | Family | Citation | STRING Data Available |
|---|---|---|---|---|
| Saccharomyces cerevisiae | Ascomycota | Saccharomycetaceae | [1] | Yes |
| Saccharomyces paradoxus | Ascomycota | Saccharomycetaceae | [2] | No |
| Saccharomyces mikatae | Ascomycota | Saccharomycetaceae | [2] | No |
| Saccharomyces bayanus | Ascomycota | Saccharomycetaceae | [3] | No |
| Saccharomyces kudriavzevii | Ascomycota | Saccharomycetaceae | [3] | No |
| Lachancea kluyverii | Ascomycota | Saccharomycetaceae | [4] | Yes |
| Naumovozyma castellii | Ascomycota | Saccharomycetaceae | [3] | No |
| Candida glabrata | Ascomycota | Saccharomycetaceae | [5] | Yes |
| Candida albicans | Ascomycota | Debaryomycetaceae | [6] | Yes |
| Candida parapsilosis | Ascomycota | Debaryomycetaceae | [7] | Yes |
| Fusarium graminearum | Ascomycota | Nectriaceae | [8] | Yes |
| Magnaporthe oryzae | Ascomycota | Magnaporthaceae | [9] | Yes |
| Aspergillus fumigatus | Ascomycota | Trichocomaceae | [10] | No |
| Aspergillus nidulans | Ascomycota | Trichocomaceae | [11] | Yes |
| Neurospora crassa | Ascomycota | Sordariaceae | [12, 13, 14] | Yes |
| Neurospora discreta | Ascomycota | Sordariaceae | [12, 13, 14] | No |
| Neurospora tetrasperma | Ascomycota | Sordariaceae | [12, 13, 14] | No |
| Cryotpococcus neoformans | Basidiomycota | Tremellaceae | [1] | No |

Table S1: Basic information on species used in analysis, including the citation corresponding to the RNA-Seq data used and whether or not STRING data was available at the time of analysis.

| Method | Correlation | TPR (S.D) | FPR (S.D) | FDR (S.D) | Accuracy (S.D) |
|---|---|---|---|---|---|
| PCM | $\rho_C$ | 0.568 (0.0003) | 0.031 (0.0037) | 0.051 (0.0056) | 0.769 (0.0018) |
| cor.test() | $\rho_U$ | 0.640 (0.0011) | 0.216 (0.0102) | 0.252 (0.0087) | 0.712 (0.00346) |
| Fraser et. al. | $\rho_C$ | 0.629 (0.0531) | 0.041 (0.0195) | 0.060 (0.0234) | 0.794 (0.0189) |
| | $\rho_U$ | 0.578 (0.0426) | 0.086 (0.0293) | 0.128 (0.0314) | 0.746 (0.0162) |
| Martin and Fraser | $\rho_C$ | 0.580 (0.0121) | 0.031 (0.0009) | 0.050 (0.0009) | 0.775 (0.0057) |
| | $\rho_U$ | 0.390 (0.0195) | 0.021 (0.0013) | 0.050 (0.0015) | 0.685 (0.0092) |

Table S2: Performance metrics for detecting coevolution using simulated data. By excluding potential false positives from the simulated binding group (i.e. STRING Score $< 400$), the overall accuracy of the methods improves, but the approaches based on the phylogenetically-corrected correlation $\rho_C$ remain superior.

#### Quantifying functional-relatedness via Gene Ontology terms

One might imagine proteins which have more overlapping GO terms are involved in more of the same functional processes, and thus would show stronger coevolution of gene expression. To quantify functional-relatedness via GO terms, the Jaccard index was used. Briefly, or a protein pair with GO terms A and B, the Jaccard Index is defined as

$$\text{Jaccard Index} = \frac{|A \cap B|}{|A \cup B|}$$

#### Simulations

All simulations were carried out using **mvMORPH**. Data were simulated from the binding group allowing evolutionary covariance term $\text{Cov}_E$ to be non-zero and simulated data from control group forcing $\text{Cov}_E = 0$. The binding group was simulated using the corresponding MLEs of the evolutionary rate matrix and ancestral state estimates from the real data. The control group was simulated similarly, but the evolutionary covariance $\text{Cov}_E$ parameter was fixed to be 0.0 (i.e. independent evolution of gene expression). Simulations used standard error estimates from the real data.

#### Supplemental Figures

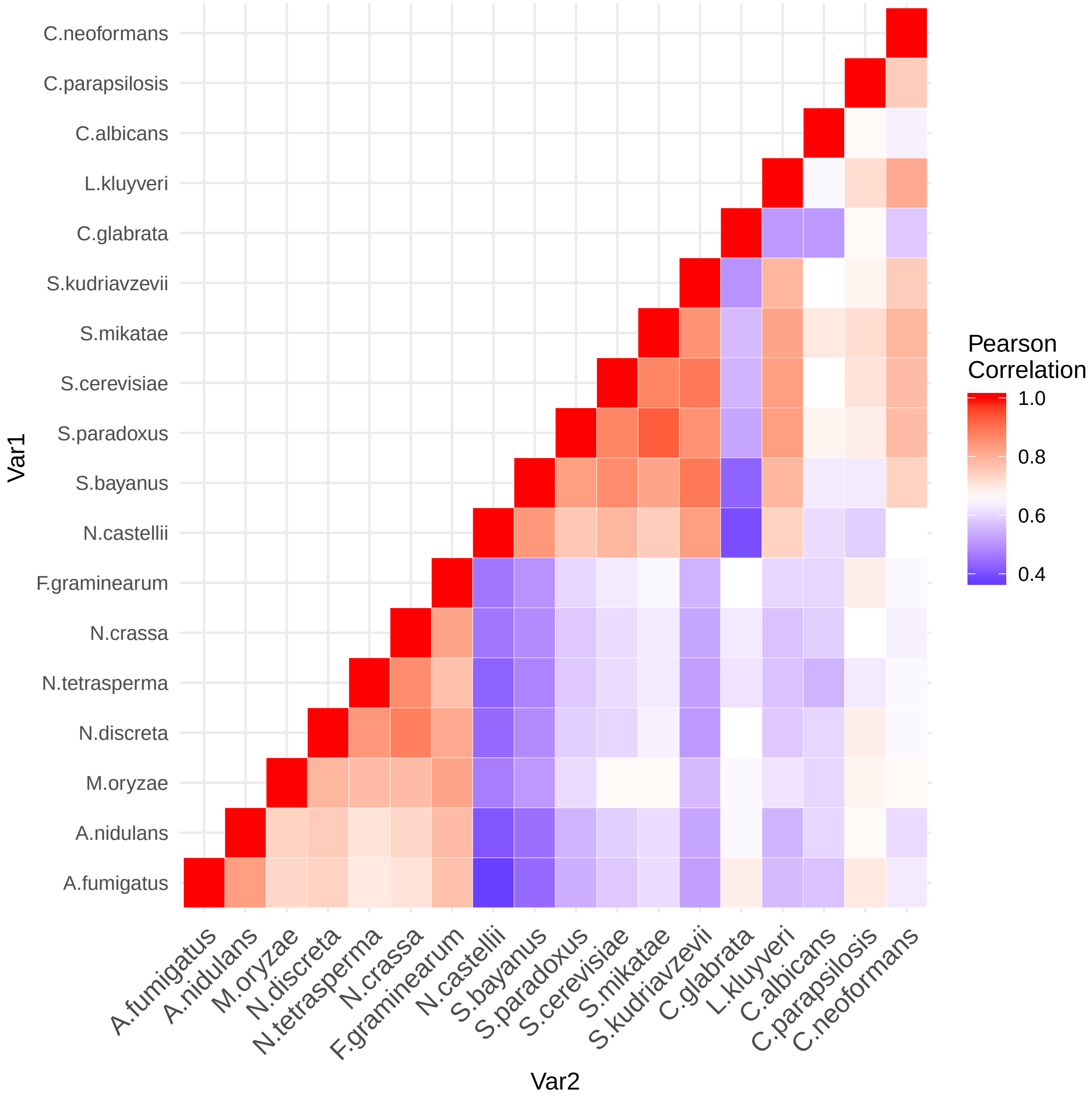


Figure S1: Heatmap demonstrating the correlation between normalized gene expression values of the 18 fungal species. Species which are more closely related tend to show higher correlations in overall gene expression patterns. *Candida* species appear to be exceptions, although gene expression is still moderately correlated with the other *Saccharomycotina* species.

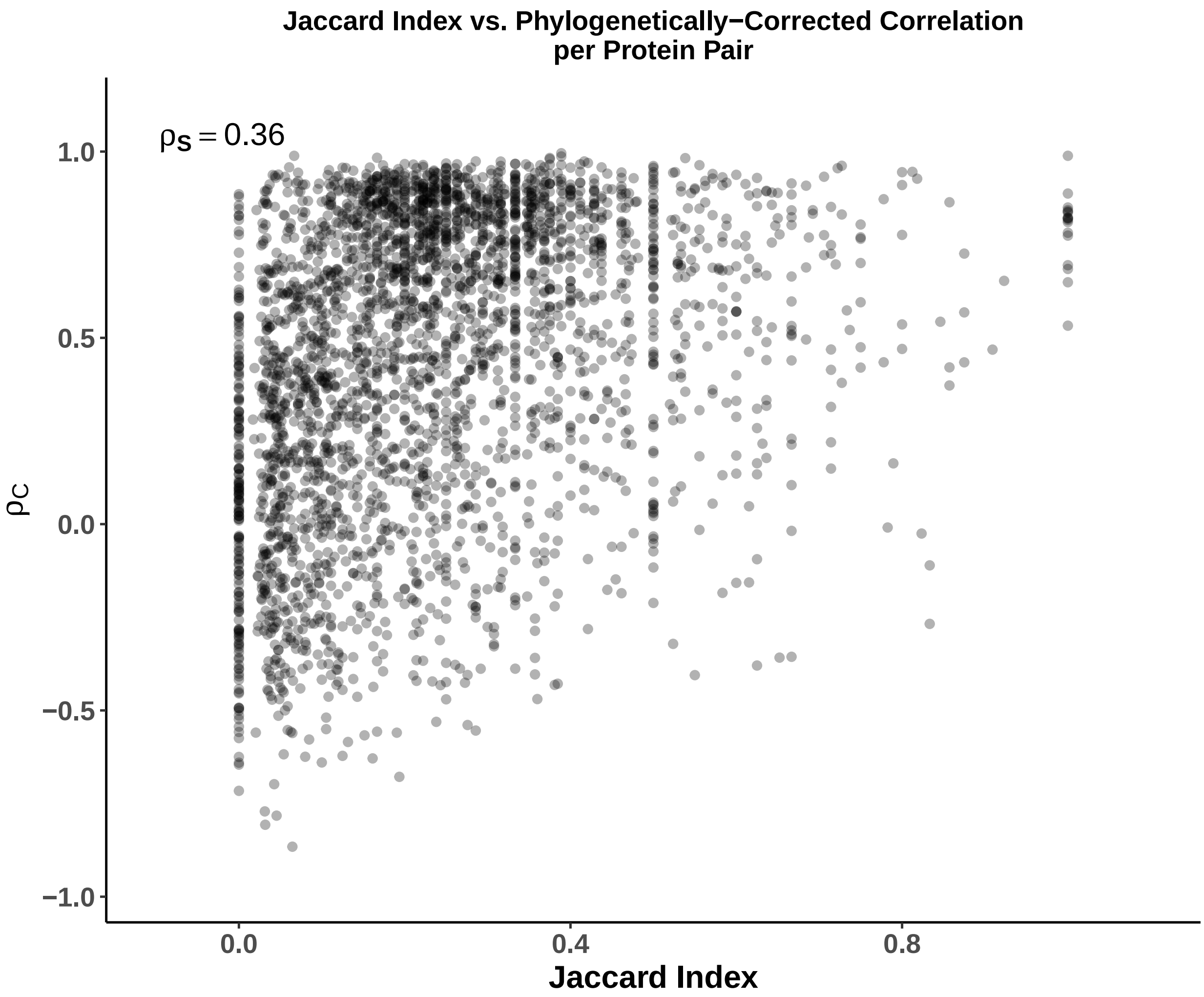


Figure S2: Pairs of proteins with more overlapping GO terms tend to show stronger coevolution of gene expresssion (Weighted Spearman Rank Correlation $\rho_S = 0.36$, 95% CI: 0.306 – 0.408, $p < 10^{-41}$). The Jaccard Index reflects functional similarity between two proteins based on GO terms.

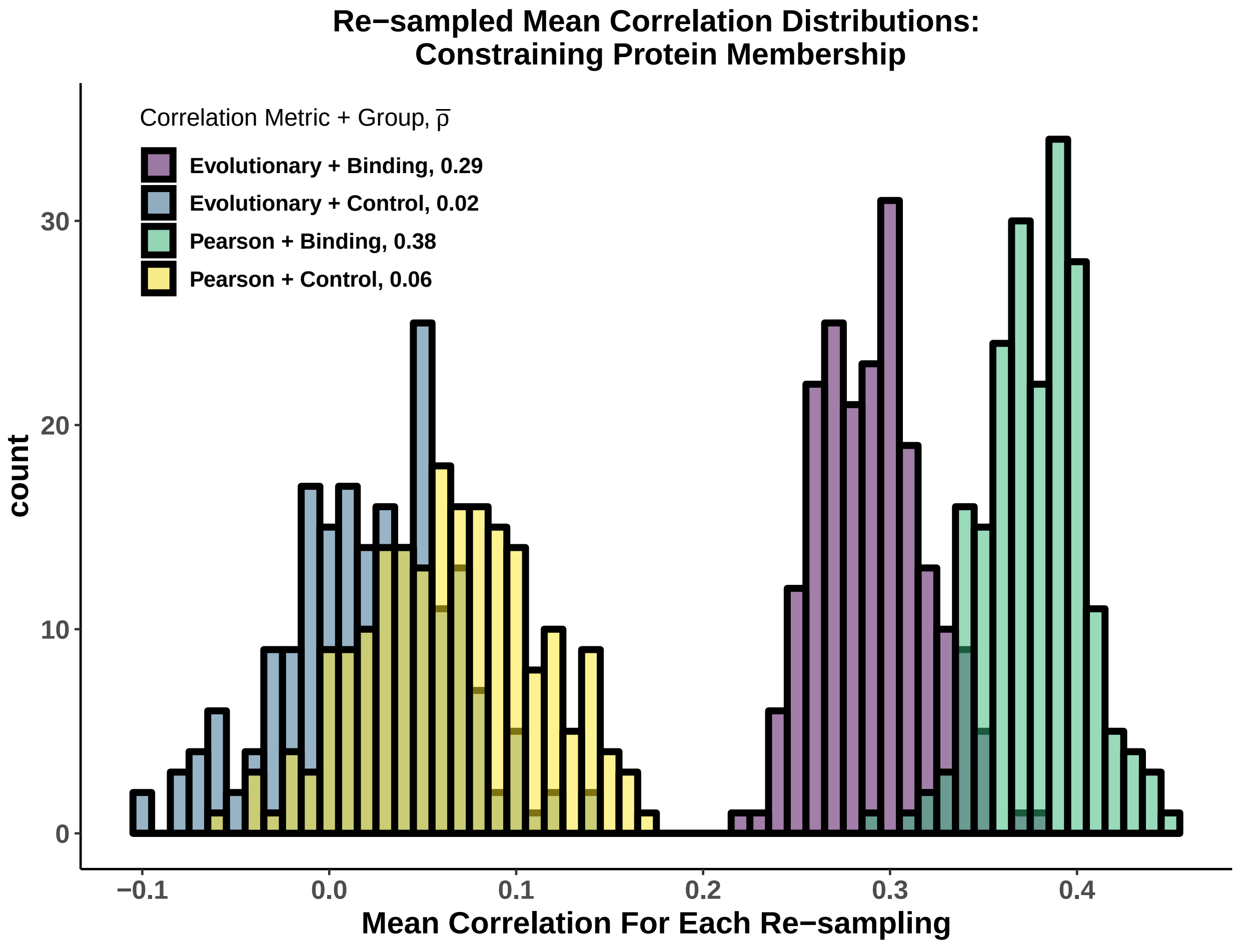


Figure S3: Distributions reflect mean phylogenetically-corrected $\bar{\rho}_C$ and phylogenetically-uncorrected $\bar{\rho}_U$ estimates for each of the 200 re-samplings of the binding and control datasets, in which each protein is restricted to being in only one pair per dataset, at max. Results are mostly consistent with results not restricting protein membership, although there does appear to be less discrepancy between the binding and control groups.

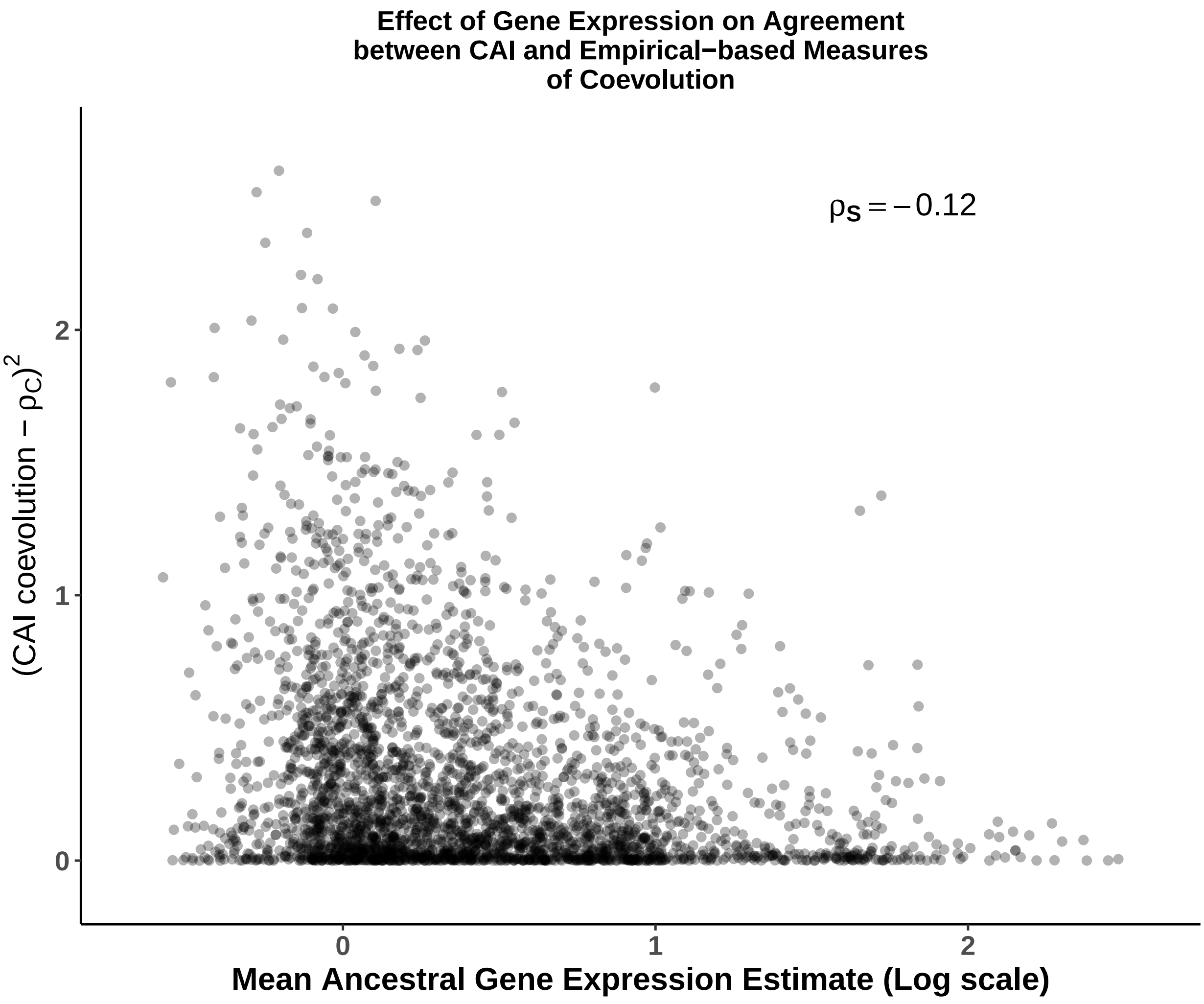


Figure S4: Determining the impact of gene expression on the agreement between CAI coevolution and $\rho_C$, as measured by the squared difference between the two metrics. A negative correlation (Weigthed Spearman Rank $\rho_S = -0.12$, $p < 10^{-4}$), indicates protein pairs which are, on average, more highly expressed tend to show less discrepancy between the CAI and empirical-based measures of gene expression coevolution.

#### Results without filtering genes violating BM assumption

The analysis was repeated not excluding genes which violated the BM assumption. This does appear to reduce some of the weighted Spearman rank correlations, but the overall conclusions are consistent with the more conservative dataset.

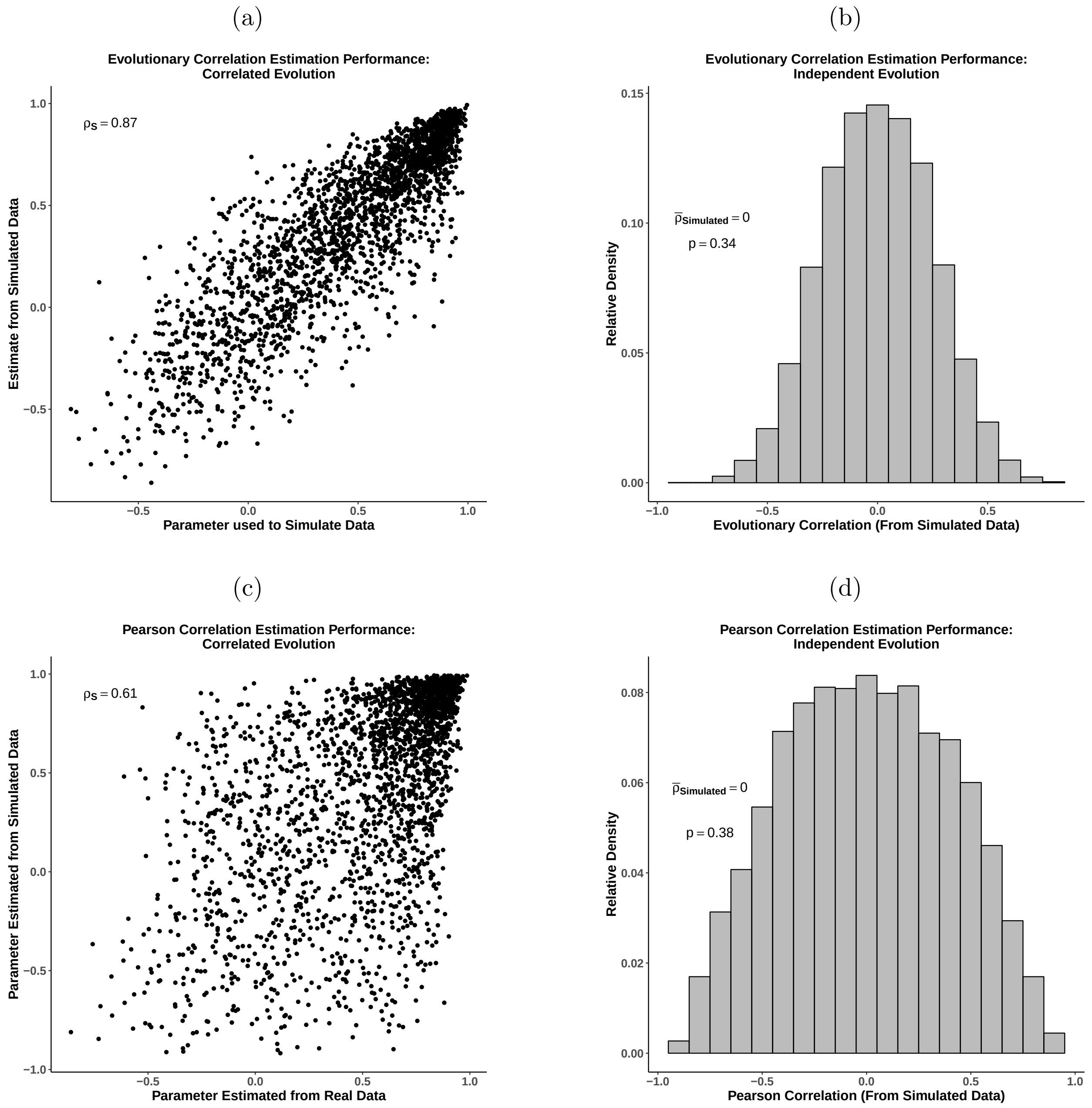


Figure S5: Estimates of phylogenetically-corrected correlation $\rho_P$ and phylogenetically-uncorrected correlation $\rho_U$ from Brownian Motion simulations when phylogenetically-corrected correlation $rho_C$ is (a,c) allowed to vary from 0.0 and (b,d) is restricted to be 0.0 (i.e. independent evolution of gene expression). (a) Comparison of $\rho_C$ from simulated data to the MLEs of $\rho_C$ used to simulate data allowing for coevolution of gene expression. Estimates from simulated data are strongly correlated with MLE from real data. (b) Distribution of $\rho_C$ estimated from simulated data forcing independent evolution of gene expression (i.e. $\rho_E = 0.0$ for all protein pairs in data set). Distribution is centered around 0.0, as expected under the null (One-Sample t-test, $p = 0.34$). (c,d) Same as (a) and (b), but for $\rho_U$. Results deviate much more from expectation when not accounting for phylogeny.

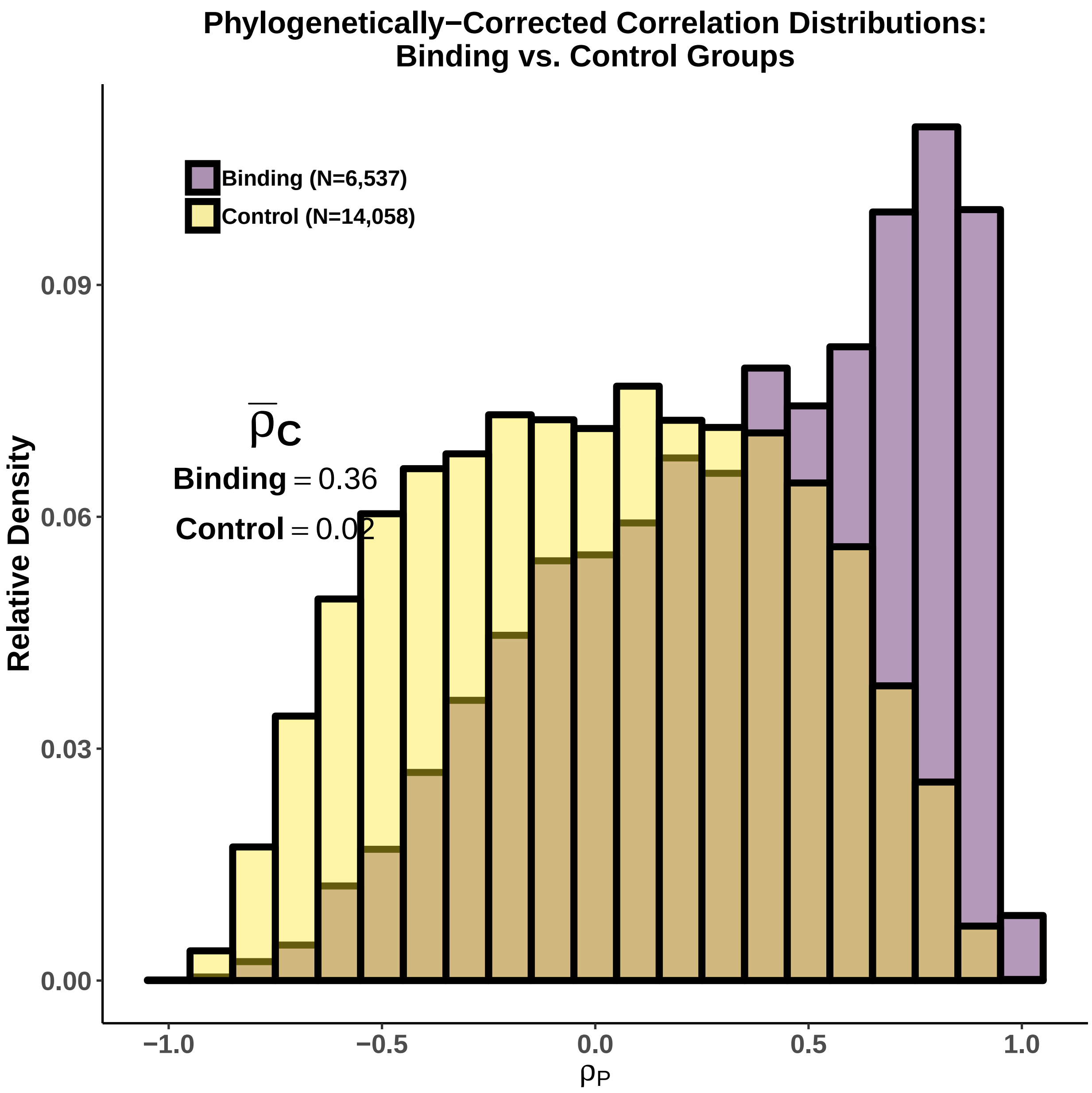


Figure S6: Comparing the distributions of the phylogenetically-corrected correlation $\rho_C$ for the binding (purple) and control (yellow) groups. Mean values for the binding and control group phylogenetically-corrected correlation $\rho_C$ distributions are 0.36 ($p < 10^{-200}$) and 0.02 ($p < 10^{-8}$), respectively.

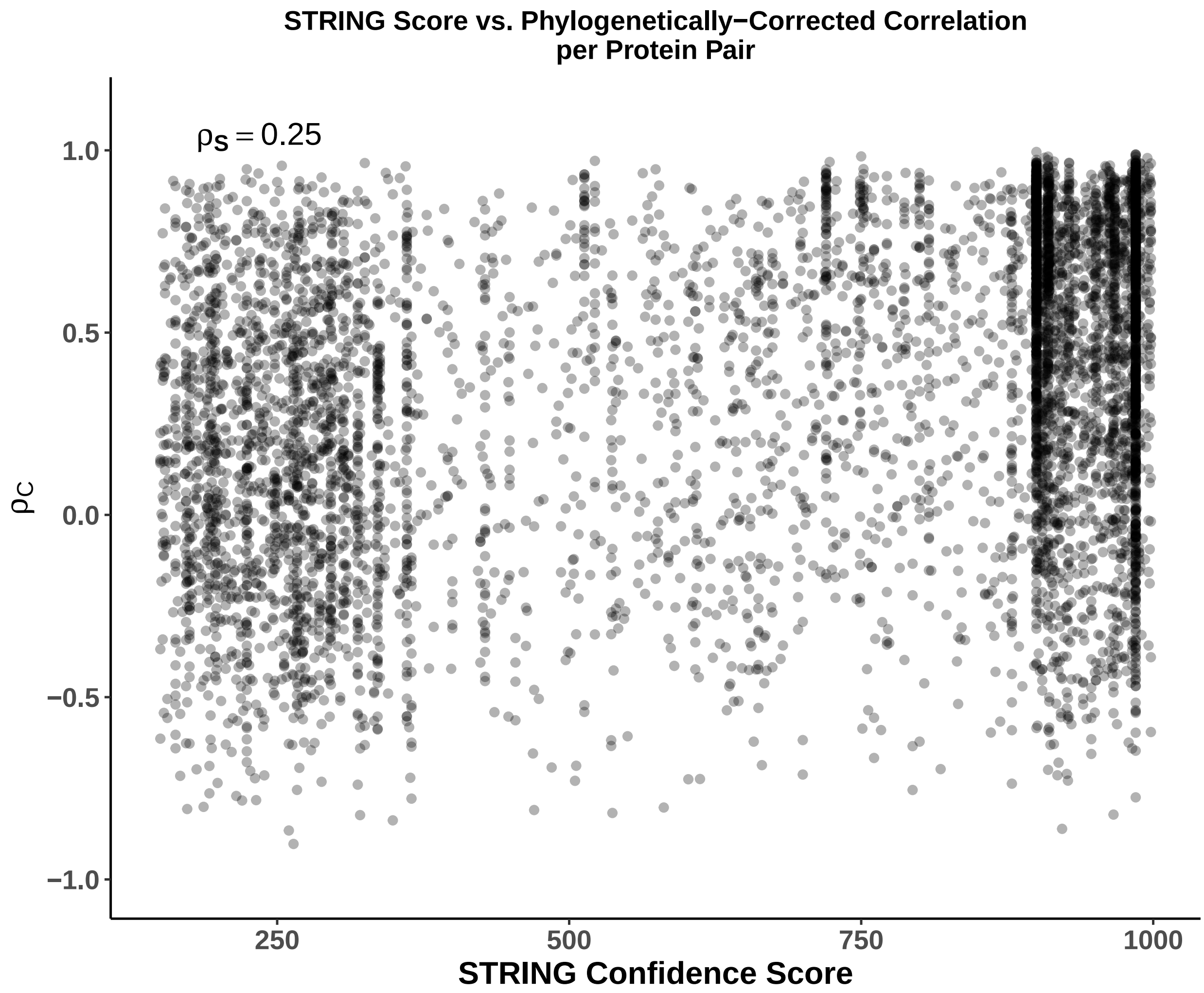


Figure S7: Effects of metric representing functional-relatedness on phylogenetically-corrected correlation $\rho_C$. Positive weighted Spearman rank correlation ($\rho_S = 0.25$, $p < 10^{-}48$) between the STRING score and phylogenetically-corrected correlation $\rho_C$ indicates more confident and/or conserved interactions tend to have higher $\rho_C$, indicating stronger coevolution at the gene expression level.

(a) (b)

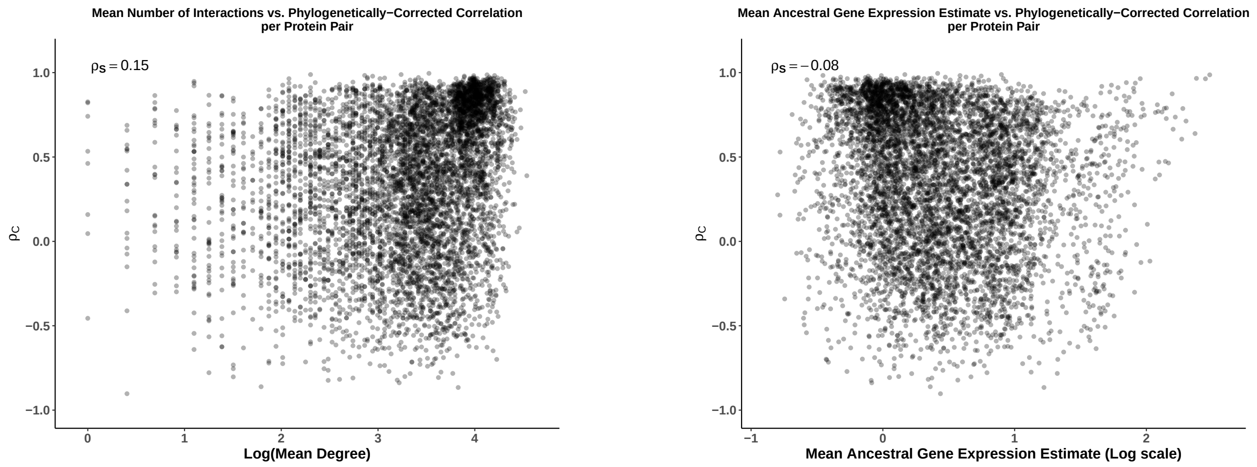


Figure S8: The relationship of (a) the mean degree (average number of interactions between a protein pair) and (b) mean ancestral gene expression estimate with the phylogenetically-corrected correlation $\rho_C$ for the binding group. Both protein pair metrics are weakly, but significantly correlated with the phylogenetically-corrected correlation $\rho_C$: weighted Spearman rank correlation $\rho_S = 0.15$ ($p < 10^{-10}$) for mean degree and $\rho_S = -0.08$ ($p < 10^{-4}$) for mean ancestral gene expression. This suggests both metrics are poor predictors of the strength of coevolution of gene expression between protein pairs.

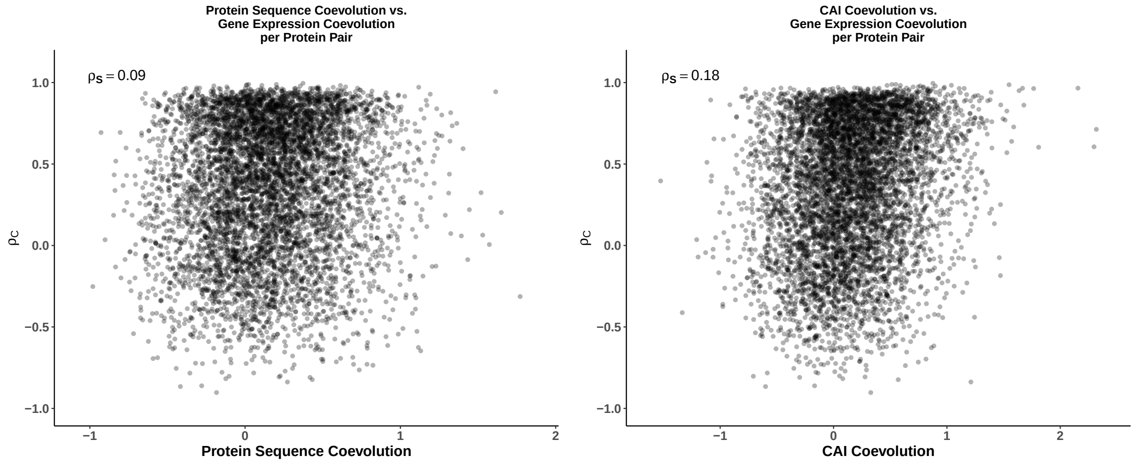


Figure S9: (a) Comparing coevolution of gene expression, represented by the phylogenetically-corrected correlation $\rho_C$, and protein sequences, as described in the main text. There is a weak but significant correlation (Weighted Spearman Rank Correlation $\rho_S = 0.09$, $p < 10^{-4}$) between the measures of gene expressions and protein sequence coevolution. (b) A similar comparison using the measures of CAI coevolution as described in main manuscript. Again, there is a weak, but significant correlation (Weighted Spearman Rank correlation $\rho_S = 0.18$, $p < 10^{-18}$).
